# Can Plitidepsin Be Used as an Antiviral Against RSV?

**DOI:** 10.1101/2025.02.27.640515

**Authors:** Charlotte Estampes, Jenna Fix, Julien Sourimant, Priscila Sutto-Ortiz, Charles-Adrien Richard, Etienne Decroly, Marie Galloux, Jean-François Eléouët

## Abstract

Human respiratory syncytial virus (HRSV) is a main cause of acute lower respiratory tract infections in infants, the elderly and the immunocompromised patients. Although vaccines have recently been approved for the elderly and for pregnant women, there is no curative treatment for HRSV. HRSV replicates in the cytoplasm of infected cells, and transcription and replication of the viral genome depend on the viral RNA polymerase complex, which recruits cellular factors for RNA synthesis. Among them, the eucaryotic translation elongation factor 1A (eEF1A) was previously shown to be critical for HRSV replication. eEF1A activity can be inhibited by plitidepsin (Aplidin), a cyclopeptide extracted from ascidian Aplidium albicans, which was shown highly potent against SARS-CoV-2, with a 50% inhibitory concentration (IC_90_) of 0.70 to 1.62 nM depending on the cell line. Here, we investigated whether plitidepsin could also inhibit HRSV replication. We found that plitidepsin inhibited HRSV replication with an IC_50_ of ≈3 nM in cell cultures. However, further investigation revealed that plitidepsin has pleiotropic effects, affecting the translation of both cellular and viral proteins in a similar manner. Overall, our results show that plitidepsin blocks cellular translation and indicate that plitidepsin induces a proteasome-mediated degradation of eEF1A, also showing the dependance of HRSV replication to cellular factors such as eEF1A. These results thus highlight an original mechanism of action of plitidepsin on eEF1A, which render the use of this compound for antiviral therapy very risky.

## Introduction

Human respiratory syncytial virus (HRSV) is the commonest cause of lower respiratory tract infection in young children worldwide and the first cause of their hospitalization (1). HRSV is estimated to infect about 33 million children leading to more than 3 million hospitalizations, and 26,000–152,000 deaths in children under 5 years, each year (2). The global healthcare costs of HRSV-associated infections in young children in 2017 were estimated to be US$5.45 billion (3). In a systemic multisite study, HRSV was shown to be the first etiological agent responsible for severe pneumonia (more than 30%) in hospitalized children in Asia and Africa (4). HRSV infections are also associated with significant morbidity and mortality in the elderly and immunocompromised people (5),(6). The true burden of disease in adults is likely significantly under-recognized, and recent studies indicate that the HRSV impact is similar to that of seasonal influenza in adults older than 65 years (7),(8),(9),(10),(11),(12).

HRSV vaccine research has been ongoing for nearly 60 years without success, and only in 2023, two vaccines based on stabilized HRSV prefusion F protein, Arexvy (GSK) and Abrysvo (Pfizer), were approved for medical use for adults aged 60 or older both in the USA and Europe. In September 2023, Abrysvo was also approved in the USA for pregnant women at 32-36 weeks’ gestation to prevent HRSV infection in newborn children. Finally, Moderna received U.S. FDA and E.U. approval for RSV Vaccine mRESVIA (13). The only pharmaceutical intervention since 1998 has been passive prophylaxis with Palivizumab, a monoclonal antibody targeting the fusion protein thereby limiting the HRSV entry. However, its use was limited to high-risk infants because of the elevated cost and moderate efficacy (14). But recently, nirsevimab (Beyfortus®), a long-acting monoclonal antibody, was approved by several regulatory agencies around the world for the prevention of HRSV infections in newborns and infants (15). A single intramuscular injection of this antibody should protect infants for an entire season compared with monthly doses, and reduced costs (vaccine-like pricing expected) allowing for administration to all infants. However, there is still no specific curative treatment against HRSV.

HRSV is a nonsegmented single-stranded negative-sense RNA virus of the *Mononegavirales* order, *Pneumoviridae* family, and *Orthopneumovirus* genus (16). The HRSV genome is approximatively 15.2 kb long, and contains 10 genes encoding 11 proteins (17). Replication and transcription rely on four of these proteins: the nucleoprotein N involved in genome and antigenome encapsidation, forming the ribonucleoprotein complex (NC), the RNA-dependent RNA polymerase L which exhibits all the enzymatic activities required for viral replication, transcription and RNA capping, its cofactor the phosphoprotein P responsible for recruitment of L on the NC template, and the transcription factor M2-1 that has been described as an “antiterminating” factor during transcription and interacts with P and viral mRNAs (18). All viral RNA synthesis take place in viral factories also called cytoplasmic inclusion bodies (IBs) (19), which concentrate the viral proteins L, N, P, and M2-1, but also cellular proteins, such as the protein phosphatase 1 (PP1), HSP70 or the human translation elongation factor eEF1A, all involved in HRVS replication (20),(21),(22). Interestingly, it was found that many RNA viruses utilize eEF1A for replication, although the mechanisms by which they do this differ (23). In a previous study in which eEF1A was knocked down or inhibited with didemnin B, it was suggested that eEF1A plays a key role in the regulation of F-actin stress fiber formation required for HRSV assembly and release, but with no effect on HRSV genome replication (24). Furthermore, eEF1A was found associated with the N and P proteins in infected cells by using a proximity ligation assay and co-immunoprecipitation (22). Recently, plitidepsin (Aplidin), an analog of didemnin B and a potent anti-cancer agent targeting eEF1A2 (K_D_=80lrnM) (25), was shown highly potent against SARS-Cov-2 by targeting eEF1A, with a 90% inhibitory concentration (IC_90_) of 0.88 nM (26). As suggested by the authors, since HRSV also uses eEF1A for viral replication, inhibition of eEF1A could be a new strategy to limit HRSV propagation/infection. In this work, we thus investigated the effect of plitidepsin on HRSV replication. We show that plitidepsin inhibits HRSV replication in infected cells as well as a minigenome reporter system with an IC_50_ of ≈3 nM in cultured cells. Further mechanistic investigation revealed that plitidepsin induces the degradation of eEF1A, inhibiting the translation of both viral and cellular proteins in a similar range of concentration.

## Methods

### Cells

BSRT7/5 cells (BHK-21 cells that constitutively express the T7 RNA polymerase) (27) and HEp-2 cells (ATCC: CCL-23) were maintained respectively in DMEM and MEM supplemented with 10% heat-inactivated fetal calf serum (FCS), with 2 mM glutamine, 100 μg/ml penicillin and 100 U/ml streptomycin. Cells were grown in an incubator at 37 °C in 5% CO_2_. Transfection were performed with 2.5µL of Lipofectamine 2000 (Thermofisher) per 1 µg of DNA according to the manufacturer instructions.

### Viruses

Recombinant human RSV rHRSV-mCherry corresponding to HRSV Long strain expressing the mCherry protein was amplified on HEp-2 cells and titrated using a plaque assay procedure as previously described (28),(29).

### Plitidepsin antiviral activity

Plitidepsin (MedChemExpress) and Carfilzomib (Cell Signaling #15022) were solubilized in DMSO at 1 mM and 5 mM as stock solutions, respectively. HEp-2 cells in 96-well plates were infected for two hours with rHRSV-mCherry at an MOI of 0.2, in the absence of FCS. The medium was then changed by the same medium with 2% FCS and containing serial dilution of plitidepsin, with a 1% final concentration of DMSO in the culture medium. At 48 h postinfection, the red fluorescence intensity of mCherry was quantified using a Tecan infinite

M200Pro spectrofluorometer with excitation and emission wavelengths of 580 nm and 620 nm, respectively. Values obtained for nontreated infected and noninfected cells were used for standardization and normalization. In parallel, the toxicity of the treatment was assessed on non-infected cells using the CellTiter-Glo Luminescent cell viability assay (Promega). The half-maximal inhibitory concentration (IC_50_) and cytotoxic concentration (CC_50_) were determined by fitting the data to the dose-response curve implemented in GraphPad version 8 software.

### Fluorescence microscopy

HEp-2 cells infected with rHRSV-mCherry were fixed with PBS-paraformaldehyde 4% for 20 min at room temperature, rinsed with PBS, and permeabilized with PBS-BSA 1% - Triton X-100 0.1% for 10 min. Nuclei were stained with Hoechst 33342 (1 µg/ml) for 5 min, washed with PBS, examined under a Nikon TE200 microscope equipped with a CoolSNAP ES2 (Photometrics) camera, and images were processed with Meta-Vue software (Molecular Devices).

### Minigenome assay

BSRT7/5 cells at 90% confluence in 96-well dishes were transfected with a plasmid mixture containing 125 ng of pM/Luc, 125 ng of pN, 125 ng of pP, 62.5 ng of pL, and 31 ng of pM2-1 as well as 31 ng of pRSV-β-Gal (Promega) to normalize transfection efficiencies as described previously (30),(31). After 6 h, the transfection mix was removed and serial dilutions of plitidepsin were added for 14 h. Transfections were done in triplicate, and each independent transfection was performed three times. Cells were harvested 24 h post-transfection, then lysed in luciferase lysis buffer (30 mM Tris pH 7.9, 10 mM MgCl_2_, 1 mM DTT, 1% Triton X-100, and 15% glycerol). The luciferase and β-galactosidase (β-Gal) activities were determined for each cell lysate with an Infinite 200 Pro (Tecan, Männedorf, Switzerland) and normalized based on the values obtained for cells treated with DMSO only.

### Fluorescence-based nucleotide-incorporation assay

Each reaction contained 0.2 µM recombinant RSV L–P, 2 µM of oligonucleotide template and 2 µM of a 5’-6-FAM primer, mixed in a buffer containing 20 mM Tris-HCl pH 7.5, 10 mM KCl, 2 mM DTT, 0.01% Triton X-100, 5% DMSO, 0.2 U/mL RNasin (Ambion), and 6 mM MgCl_2_. Reactions were started by adding specific NTPs at 100 µM and incubated for 2h at 30°C. Reactions were quenched by adding an equal volume of gel-loading buffer. Samples were denatured at 70°C for 10 minutes and run on a 20% polyacrylamide urea sequencing gel for 2.5 hours at 45 W. The gel scanned on a Typhoon imager (GE Healthcare). (Unpublished methodology Sutto et al. 2024)

### Western blotting

Cells were lysed in 2X Laemmli buffer, run on a 12% polyacrylamide gel and transferred to a nitrocellulose membrane using the Trans-Blot Turbo system (Bio-Rad). The blots were blocked with 5% nonfat milk in PBS Tween20 0.2%, and probed with either a polyclonal rabbit anti-N serum (32), a rabbit anti-GFP (Invitrogen, Whaltham, MA, USA), a mouse monoclonal anti-EF1A antibody (Santa Cruz G-8, sc377439), a mouse monoclonal anti-GAPDH antibody (Sigma-Aldrich MAB374) or a mouse anti-alpha-tubulin (DM1A, Sigma) and further incubated with HRP-coupled anti-mouse or anti-rabbit antibodies (Life Science). Membranes were revealed with the Clarity Western ECL kit (Bio-Rad) and analyzed with a ChemiDoc Touch Imaging System and Image Lab software (Bio-Rad).

### Pulse labelling of newly synthesized proteins with L-azidohomoalanine (AHA)

HEp-2 cells were infected with rHRSV-mCherry at MOI = 1 in 6-wells plates (35 mm in diameter) and incubated in MEM supplemented with 2% heat-inactivated FCS, 2 mM glutamine, 100 μg/ml penicillin and 100 U/ml streptomycin. 24 h later, serial dilutions of plitidepsin or DMSO were added to the cells for 2 h. Cells were washed twice with methionine-free (Met-) MEM (Gibco), then live-labeled (pulsed) at 37^lr^C for 4 h with 50 μM Click-iT L-azidohomoalanine (AHA; Jena Bioscience) in Met-MEM supplemented with 5–10% dialyzed FBS (Gibco) and plitidepsin. Cells were lysed in 1 ml of in RIPA buffer (50 mM Tris– HCl [pH 8.0], 150 mM NaCl, 1% Triton X-100, 2 mM, DTT, 10 mM of freshly prepared iodoacetamide [Merck] and complete protease inhibitor cocktail [Roche]). After centrifugation for 10 min at 10,000 rpm at 4°C, the supernatants were mixed with 5 μM AFDye 488-DBCO (Click Chemistry Tools, Jena Bioscience) and incubated for 2 h at 21°C. Proteins were immunoprecipitated with specific antibodies coupled to protein A sepharose beads (GE Healthcare) and analyzed by SDS-PAGE, together with the supernatants. Fluorescence was analyzed with a Fuji FLA-3000 scanner and quantified using the AIDA or Fiji softwares.

## RESULTS

### Antiviral potency of plitidepsin against HRSV

By using an siRNA approach, it was demonstrated ten years ago that downregulation of eEF1A correlates with a reduction in the amount of infectious HRSV release, accompanied by a reduction in viral genome expression, but not mRNA transcription or protein expression (22). We therefore tested whether plitidepsin, known to inhibit the eEF1A activity, displays antiviral activity against HRSV. For this purpose, HEp-2 cells were infected with rHRSV-mCherry at MOI = 0.2 in a culture medium containing increasing concentration of plitidepsin. Viral replication was quantified by mCherry fluorescence measurement on live cells 48 h post infection using the Tecan spectrofluorometer. Values were standardized and normalized by those of obtained for nontreated infected and noninfected cells. The IC_50_ was determined by dose-response curve fitting. In parallel, the toxicity of plitidepsin was evaluated on non-infected HEp-2 cells. As shown in Figure 1A, plitidepsin inhibited rHRSV-mCherry replication in a dose-dependent manner, with an IC_50_ of 3.5 nM. These data were confirmed by direct observation of mCherry fluorescence and nuclei staining on fixed rHRSV-mCherry-infected cells (Figure 1B). Noteworthy, a cytotoxic activity of plitidepsin was observed, with a calculated half-maximal cytotoxic concentration (CC_50_) close to 145 nM (Figure 1A). With a calculated selective index >30, plitidepsin thus appeared to be a potent inhibitor of HRSV replication in HEp-2 cells.

**Figure 1.**
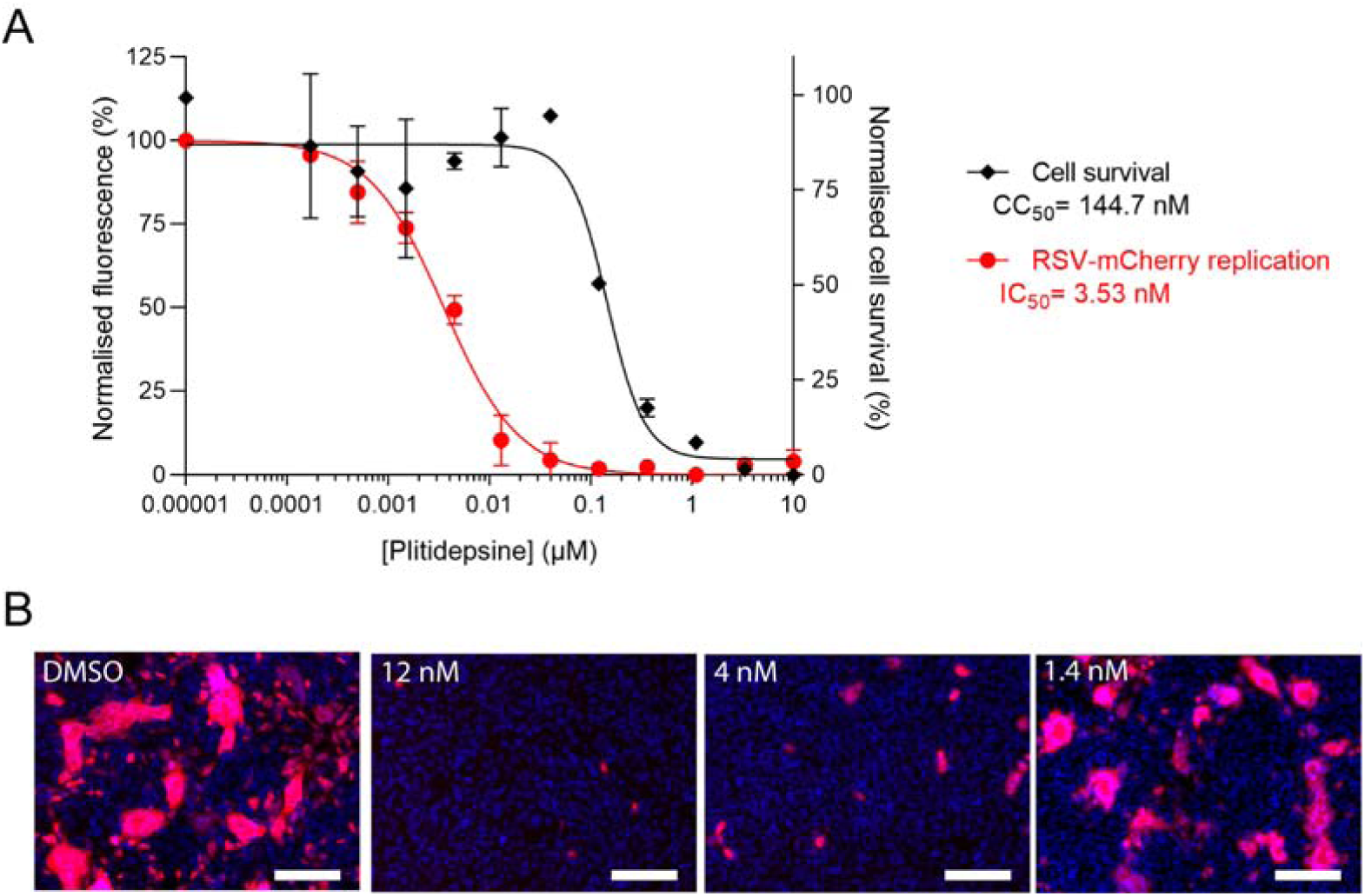
Impact of plitidepsin treatment on HRSV replication on HEp-2 cells. **(A)** Cells were infected for 2 h with rHRSV-mCherry at MOI 0.2 and the medium was then replaced to incubate cells in the presence of serial dilutions of plitidepsin for 48 h (red curve). The viral replication was quantified by measurement of the mCherry fluorescence. In parallel, cell viability upon treatment with plitidepsin was quantified in non-infected HEp-2 cells (black curve). Error bars are standard deviations from duplicates. Data are representative of three experiments. The curves were fitted in Graph Pad 8 software using a four parameters logistic (4PL) regression. Both IC_50_ and CC_50_ are indicated. **(B)** Representative images of HEp-2 cell cultures infected with rHRSV-mCherry at MOI 0.2, 48 hours post-infection in the absence or presence of plitidepsin at 12, 4, and 1.4 nM. Nuclei were colored with Hoechst 33432. Scale bars, 250 µm.

### Effect of plitidepsin on a HRSV minigenome

To determine whether plitidepsin could affect the RNA polymerase replication/transcription machinery of HRSV, we used a well-established HRSV-specific minigenome assay system (30). The pM/Luc plasmid, which contains the authentic M/SH gene junction and the Luc reporter gene downstream of the gene start sequence inserted in this gene junction, was co-transfected in BSR-T7/5 cells together with p-β-gal, pL, pP, PN, and pM2-1. Luciferase activity was determined and normalized based on the signal obtained for cells incubated in the similar media with 1% of DMSO. In parallel, β-galactosidase (β-Gal) activity was determined for each cell lysate in order to normalize transfection efficiency. Figure 2A shows that plitidepsin inhibits the function of the HRSV polymerase complex in a dose-dependent manner with an IC_50_ of ∼5 nM while showing no toxicity towards BSR-T7/5 cells. However, plitidepsin also inhibited, to a lesser extent, the β-Gal activity, the effect being visible from 3.3 nM of plitidepsin.

**Figure 2.**
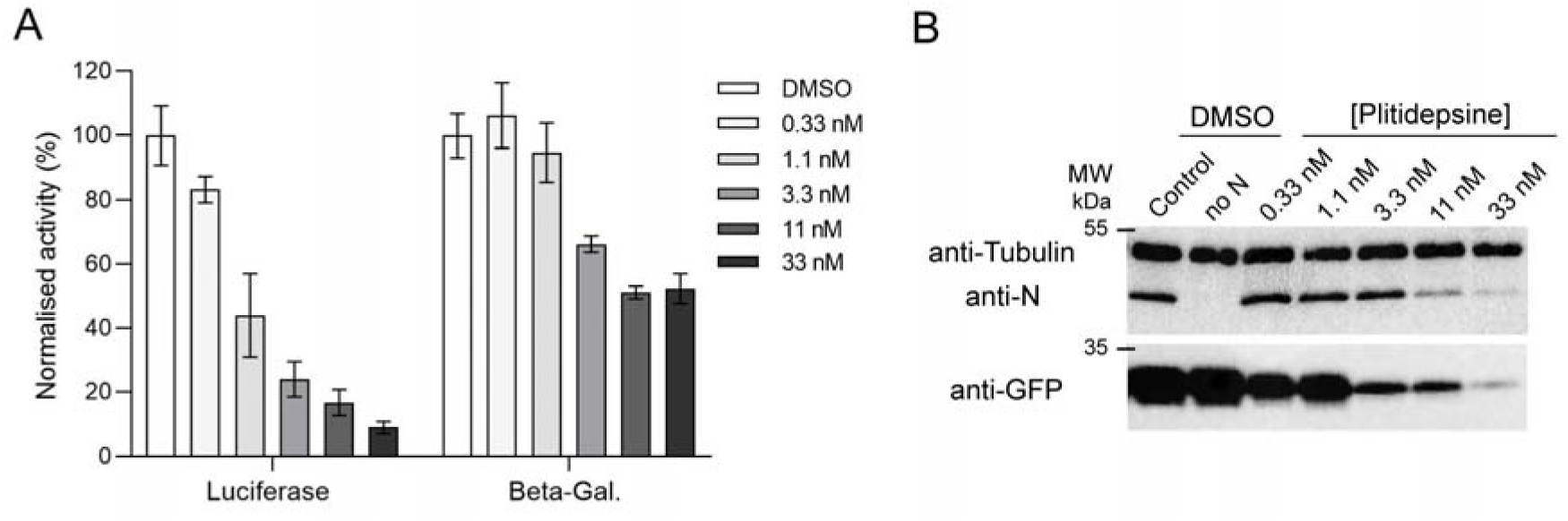
Plitidepsin effect on a HRSV minigenome. (A) BSRT7/5 cells were transfected with plasmids encoding the N, P, M2–1, and L proteins; the pMT/Luc minigenome, together with pCMV-beta-gal for transfection standardization. Serial dilutions of plitidepsin were added 6 h post-transfection after removing transfection reagents. Viral RNA synthesis was quantified by measuring the luciferase activity after cell lysis 24 h post-transfection. Each luciferase minigenome activity value was normalized based on DMSO treated cells and is the average of three independent experiments performed in triplicate. Error bars represent standard deviations calculated based on three independent experiments made in triplicate. (B) BSRT7 cells were transfected with the complete minigenome system (see above) or with the pEGFP plasmid coding for EGFP. Six hours later, the transfection mix was removed and medium containing plitidepsin at different concentrations was added. Then, 24h post-transfection, cells were lysed and analyzed by Western blotting using anti-N, anti-GFP or anti-alpha tubulin antibodies.

Since plitidepsin targets the eEF1A cellular translation factor, the inhibition of HRSV RNA polymerase activity could result from either the direct inhibition of polymerase activity of the HRSV L protein or from the inhibition of translation of the viral mRNAs coding for the viral proteins. To assess whether the plitidepsin treatment had a specific or global impact on proteins expression, cells were co-transfected either with the minigenome system or with a plasmid encoding EGFP, in the presence of increasing concentrations of plitidepsin. The expression of proteins was analyzed by Western blot. As shown on Figure 2B, a similar decrease in N and EGFP expression was observed in the presence of plitidepsin, indicating that plitidepsin had a general inhibitory effect on viral or non-viral proteins expression. In contrast, the signal corresponding to alpha-tubulin did not decrease significantly, which can easily be explained by the half-life of this cellular protein (∼8 days) (33).

### Effect of plitidepsin on *in vitro* HRSV RdRp activity

To test whether plitidepsin could also affect directly the HRSV RNA polymerase activity, we performed an *in vitro* RNA polymerase assay using a recombinant L-P complex (34),(35),(36) in the presence of increasing plitidepsin concentrations. As shown on Figure 3, no inhibitory effect of plitidepsin on the RNA synthesis was observed, even for a concentration of 100 µM of plitidepsin. These results strongly suggest that the effect of plitidepsin on viral replication was not due to a direct effect of the RdRp complex but mainly due to its impact on protein expression.

**Figure 3.**
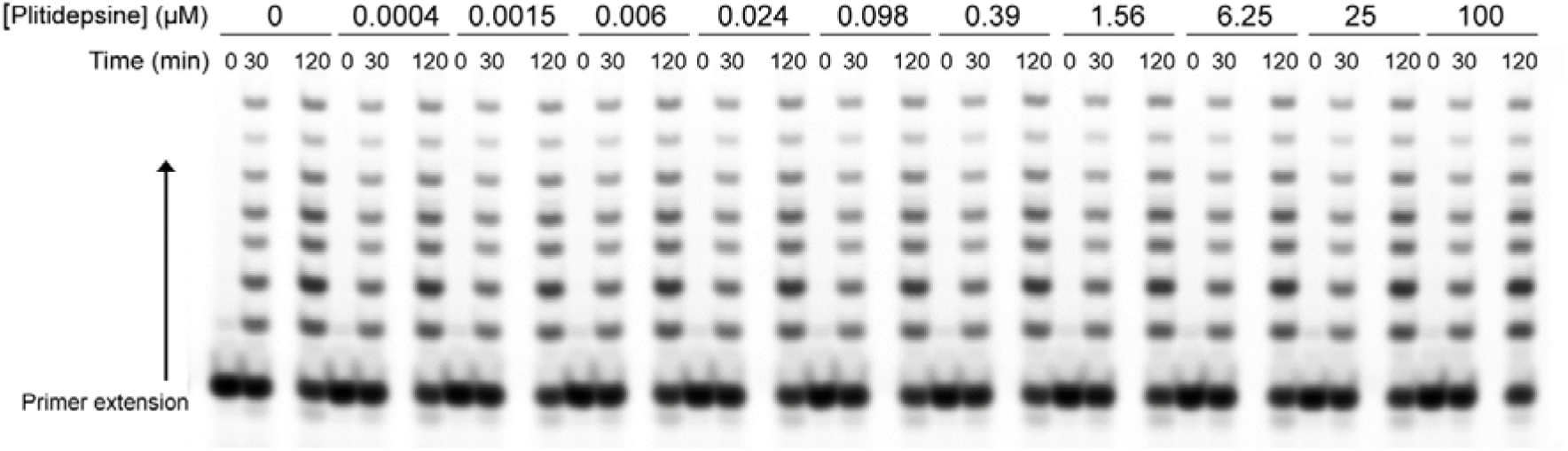
*In vitro* effect of plitidepsin on HRSV RNA polymerase (L-P complex) activity. Primer extension polymerase assay was performed in the presence of increasing concentration of plitidepsin. Briefly, 200 nM of RSV L/P was incubated with 2 μM of 11 nucleotide length RNA template and 4 nucleotide primer in presence of increasing concentrations of plitidepsin. The reactions were incubated for 30’ and 120 min at 30°C and the reaction products were separated on a 20 % UREA-PAGE gel before gel scanning on a Typhoon imager.

### Effect of plitidepsin on cellular and viral proteins neosynthesis

To further investigate the impact of plitidepsin on viral and cellular RNA translation, we used a complementary approach, consisting in a pulse labelling of newly synthesized proteins with the methionine analog L-azidohomoalanine (AHA) revealed by the fluorescent dye AFDye 488-DBCO based on Click chemistry (37). At 24 h postinfection, HEp-2 cells were incubated with serial dilutions of plitidepsin and then labelled with AHA for 1 h. Then, cells were lysed and the HRSV N protein or the cellular α-tubulin protein were immunoprecipitated. Immunoprecipitated proteins and total cell lysates were analyzed by SDS-PAGE and the fluorescence was quantified. As shown on Figure 4A and 4B, a similar drop in protein synthesis was observed for N and α-tubulin, but also for the total proteins present in the cell lysates for concentrations ≥ 15 nM.

**Figure 4.**
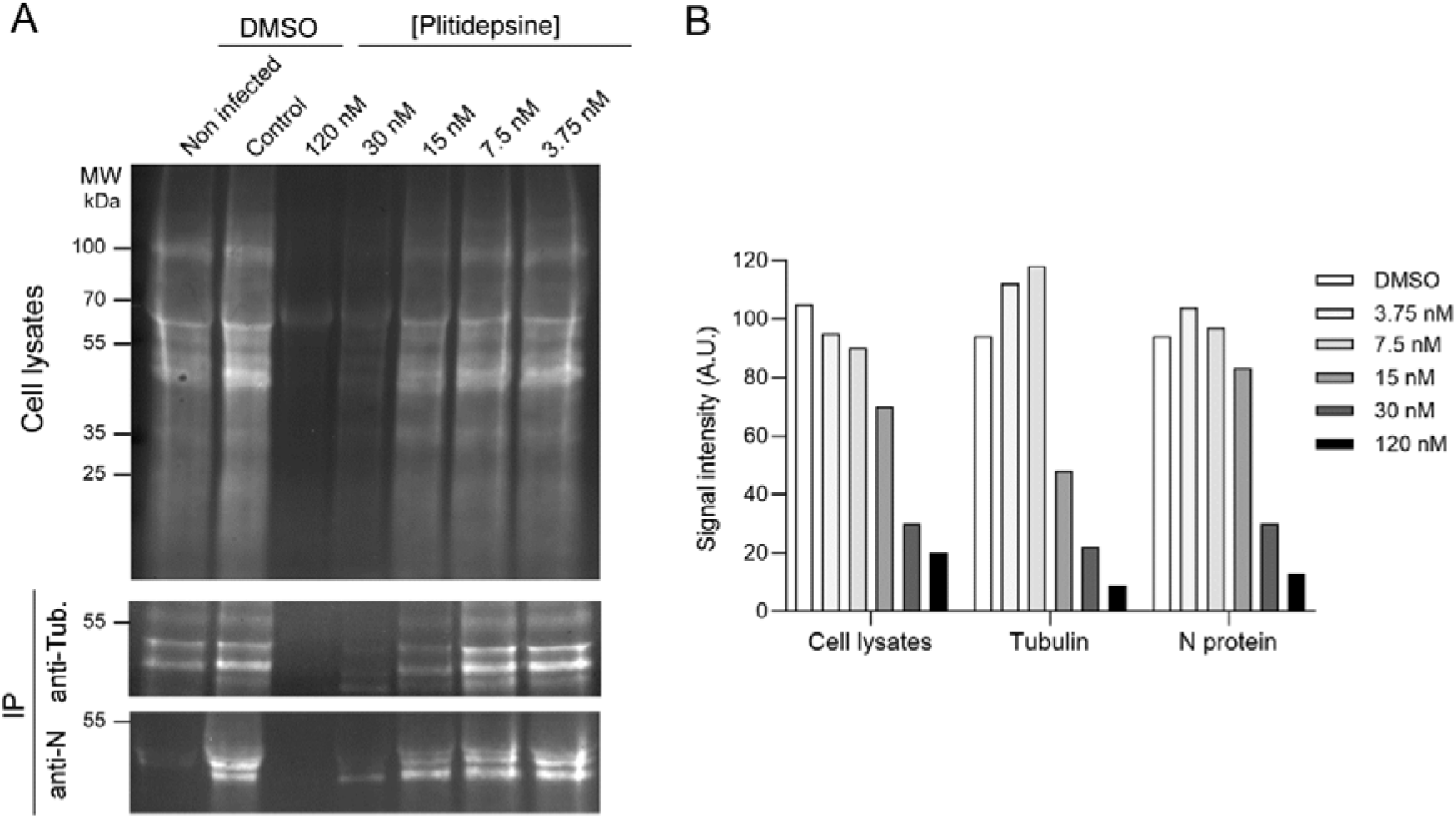
Compared effect of plitidepsin on viral or cellular proteins synthesis. HEp-2 cells in 6 well plates were infected with rHRSV-mCherry at MOI = 1. 24 h later, medium was changed for protein labelling with AHA with serial dilutions of plitidepsin. After 4 h, cells were lysed and AHA-labelled proteins were revealed with AFDye 488-DBCO, then immunoprecipitated with anti-N or anti-alpha-tubulin antibodies. Immunoprecipitated proteins or supernatants were resolved by SDS-PAGE (A) and fluorescence in the gels was measured using a Fuji FLA-3000 scanner (B).

### Plitidepsin induces proteasome-mediated degradation of eEF1A

The mechanism by which plitidepsin inhibits the activity of the host factor eEF1A is still debated. It was shown that overexpression of a negative dominant mutant an Ala399 → Val (A399V) of eEF1A reduced sensitivity of cancer cells to didemnin B (38) as well as SARS-CoV-2 to plitidepsin by a factor >10 (26). Interestingly, this A399V substitution also confers resistance to the structurally unrelated ternatin-4 (38), suggesting that these molecules interact with the same binding site on eEF1A (26). However, the mechanism of action of ternatin-4 was recently studied in more details and revealed that it was mediated by ubiquitination and proteasome degradation of eEF1A (39). We thus wondered whether plitidepsin could also induce degradation of eEF1A by a similar mechanism. BSRT7 cells were treated with serial dilutions of plitidepsin in the presence or absence of the proteasome inhibitor carfilzomib and the expression of eEF1A or the control GAPDH were analyzed by Western blot. As shown on Figure 5, at the RSV inhibitory concentration of plitidepsin, minigenome activity or translation inhibition, eEF1A expression was impaired. However, expression of eEF1A was restored upon treatment in the presence of the proteasome inhibitor Carfilzomib. These results thus strongly suggest that plitidepsin induces the degradation of eEF1A by ubiquitination and proteasome degradation in a similar way to ternatin-4.

**Fig. 5.**
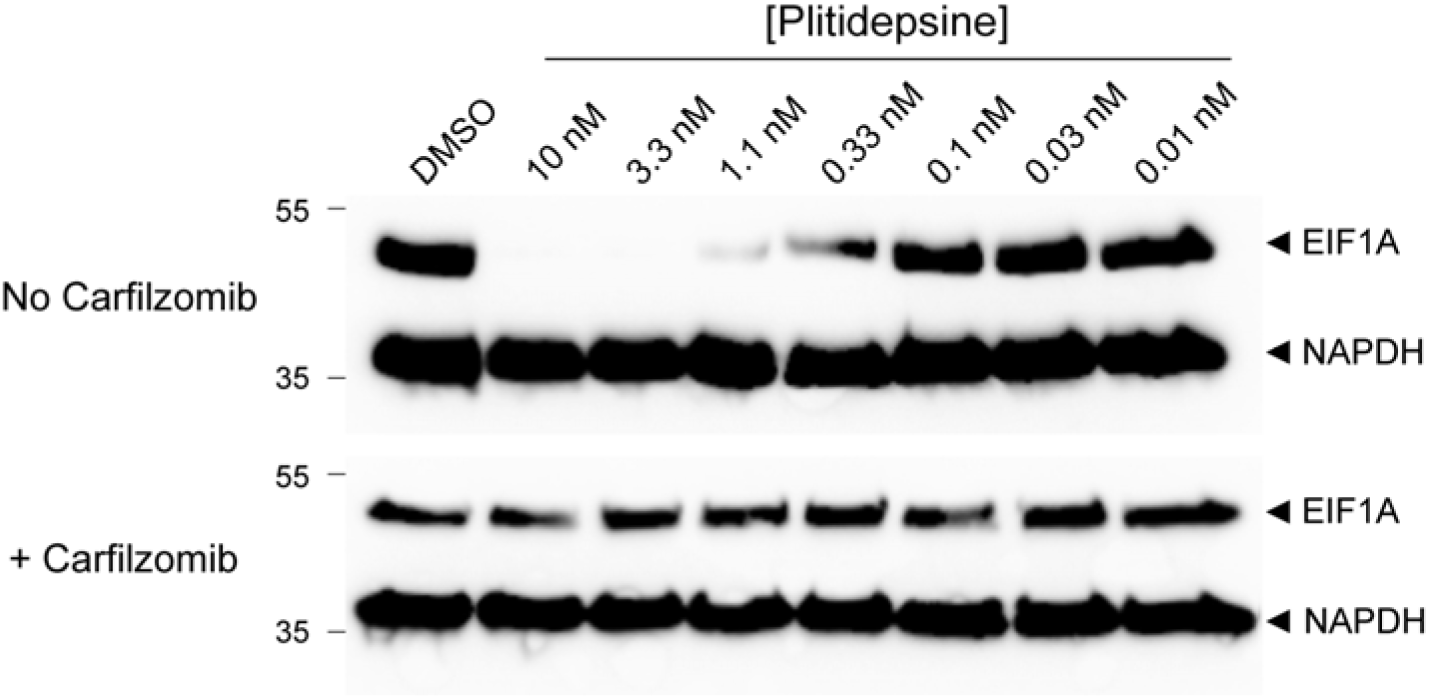
Antiviral mechanism of action of plitidepsin is mediated through proteasome-mediated degradation of eEF1A. BSRT7/5 cells were treated with serial dilutions of plitidepsin or DMSO for 18H in the presence or not of the proteasome inhibitor Carfilzomib (500 nM) and the expression of eEF1A or GAPDH were analyzed by WB.

## DISCUSSION

The mammalian translation elongation factor eEF1A is an essential GTPase and the second most abundant intra-cellular protein after actin (3% of the total cellular protein). It is localized extensively in the cytoplasm and nucleus (40),(41),(23). The canonical function of eEF1A is to deliver amino acyl tRNAs to the ribosomal A site during the elongation stage of protein synthesis. In addition to its canonical functions in transporting aa-tRNA to the ribosome, eEF1A is found to be involved in cellular mechanisms such as regulation of cytoskeleton organization by interacting with actin and tubulin, protein degradation mediated by the proteasome, nuclear aa-tRNAs and protein export, signaling transduction pathway concerning apoptosis and oncogenesis, or binding to viral RNA (42),(41),(43). There are two variants of eEF1A, eEF1A1 and eEF1A2, that share 92% amino acid identity (44). In contrast to the ubiquitous expression of eEF1A1 in many cell types, eEF1A2 expression is limited to the terminally differentiated cells of the brain, heart, and skeletal muscle (45). The two eEF1A variants have similar translation activity but may differ with respect to their secondary, “moonlighting” functions. eEF1A1 also plays an important role in the process of heat shock stress response. eEF1A2 activates Akt in a PI3K-dependent fashion, stimulating cell migration, actin remodeling and invasion, and inhibiting apoptosis (23).

Like for many viruses (23),(41), eEF1A has been identified as an important cellular factor for HRSV replication (22). As an abundant, multifunctional protein, it is not surprising that many viruses have adapted to use eEF1A as a cofactor for viral transcription, translation, assembly, and pathogenesis. So, eEF1A was shown to bind to some viral RNA structures, some viral structural and nonstructural proteins, or to interact with viral polymerase complexes, such as the one of Vesicular Stomatitis Virus (VSV) (46),(41). Like for HRSV, VSV forms two different viral RNA polymerase complexes in infected cells: the transcriptase and the replicase. The transcriptase complex synthesizes capped mRNAs, whereas the replicase complex initiates genomic minus-strand RNA synthesis at the precise 3’ ends of the plus-strand antigenomic and negative-strand genomic RNAs. The transcriptase was described as a protein complex containing the L and P proteins, the cellular eEF1A and heat shock protein 60, and a submolar amount of cellular mRNA cap guanylyltransferase (47). The replicase complex was described as containing the viral proteins L, P, and nucleocapsid (N) but not eEF1A, heat shock protein 60, or the guanylyltransferase. Hence, it was proposed that eEF1A is important for transcription of viral mRNAs but not for genomic RNA replication in VSV.

Disentangling the canonical and noncanonical roles of eEF1A and the extent to which each contributes to viral function is essential if eEF1A is to be targeted therapeutically. It was previously shown that didemnin B treatment, a drug targeting eEF1A, protected cells from HRSV-induced cell death (22). Didemnin B did not significantly affect HRSV transcription and replication, especially at 24 h post-infection, but significantly reduced infectious virus production and release, possibly as a consequence of changes in actin stress fiber formation (24). More recently, plitidepsin (Aplidin), an analog of didemnin B that was approved in Australia for the treatment of myeloma (48), was shown to be very efficient against SARS-CoV-2 in cultured cells as well as in a mice model (26). Since it has entered clinical trials as an anti-SARS-CoV-2 drug and was shown as a favorable long-term safety profile in adult patients hospitalized for COVID-19 (49),(50),(51),(52), we wondered whether this compound could also be efficient against HRSV. Plitidepsin is a cyclic depsipeptide that was first isolated from a Mediterranean marine tunicate (*Aplidium albicans*) and, at present, is manufactured by total synthesis and commercialized by PharmaMar, S.A., as Aplidin®(53). Plitidepsin has antitumoral, and immunosuppressive activities (54). Although not fully clarified, the molecular mechanisms of action of plitidepsin against tumor cells and SARS-CoV-2 have been investigated. Plitidepsin induces cell-cycle arrest and apoptosis (55). These effects rely on the induction of early oxidative stress, the rapid activation of Rac1 GTPase, and activation of c-Jun N-terminal kinase (JNK), ERK and p38 mitogen-activated protein kinases (p38/MAPK), which finally result in caspase-dependent apoptosis (56),(57). It was determined that eEF1A is the primary target of plitidepsin, which can bind to eEF1A at the interface between domains 1 and 2 of this protein in the GTP conformation with a measured KD of 80 nM (25). However, it was proposed that the antitumoral effect of plitidepsin is not due to translation inhibition but inhibition of eEF1A binding to double-stranded RNA-dependent protein kinase (PKR). In the presence of plitidepsin, PKR would disengage from eEF1A, thereby regaining its kinase activity to initiate extrinsic apoptosis through activation of MAPK and NF-κB signaling cascades (58). Plitidepsin was also reported as an ER stress inducer by activating the unfolded protein response (59). In parallel, plitidepsin was also shown to induce the phosphorylation of eIF2α, resulting in the arrest of protein synthesis at the initiation step (59), (60). For SARS-CoV-2, it has been determined that the effect of plitidepsin is also mediated by eEF1A by using a mutated (A399V) version of eEF1A1 in 293T cells (26). It is likely that this effect is mediated by inhibition of translation of the viral proteins (26), (61),(55). Very recently, the mechanism of SARS-CoV-2 inhibition was revisited (62). In this paper, using Vero E6 cells, the authors found that plitidepsin at 50 nM reduced the translation of RNAs, including cellular RNAs, but with a higher impact on viral mRNAs translation and without affecting cellular viability. The molecular mechanisms involved cell’s proteostatic response to eEF1A blockade, namely a shift from cap or internal ribosome entry sites (IRES)-mediated translation towards a N-6-methyladenosine (m6A)-dependent translation, that could explain cell survival. They also tested plitidepsin against several other viruses including HRSV, and found an IC50 of 27 nM in Vero E6 cells, about ten times more than what we found using HEp-2 cells. For *in vivo* effects, it was also shown that treatment of a monocyte-derived macrophage cell line by plitidepsin in the same range than ours (2.5-10 nM) reduced the production of the proinflammatory cytokine IL-6 in the presence of SARS-CoV-2 virions, and that was associated with a reduction of NF-κB p65 subunit phosphorylation and of its transcription activity in the inflammatory cascade (63). This effect was also observed *in vivo* in SARS-CoV-2 or Influenza virus infected mice.

In our experiments, we observed that plitidepsin had an inhibitory effect on viral replication and minigenome activity with an IEC_50_ ≈ 3-5 nM, and a toxic effect on cells with an IC_50_ ≈ 150 nM. However, further investigation revealed that this inhibitory effect was due to a general inhibition of translation in the host cell, with no significant difference between cellular and viral proteins expression. We then wondered whether eEF1A, which is essential for cellular translation, could be actively degraded by plitidepsin treatment, as it was observed recently with ternatin-4, another eEF1A-targeting drug (39). We found that eEF1A was degraded after plitidepsin treatment for 18H at concentrations as low as 1 nM, which was inhibited by proteasome inhibitor carfilzomib, indicating that a eEF1A ubiquitination mechanism occurred, that remain to be determined. During this study it was published by Molina et al. that cellular cap-dependent translation was inhibited by plitidepsin (see above). However, using pulse-chase experiments, we found that cellular translation was mainly affected (see Figure 4).

In conclusion, we show here for the first time that treatment of cells with plitidepsin induces the degradation of eEF1A by the proteasome pathway, which is correlated with a global extinction of translation. These experiments also show the dependance of HRSV replication to the cellular factor eEF1A. Although effective concentrations of plitidepsin against HRSV are in the same range that those found for SARS-CoV-2, the side effects of plitidepsin on cells and its potential toxicity raises the question of its use *in vivo* for antiviral assays against HRSV.

## References

1. Langedijk AC, Bont LJ. 2023. Respiratory syncytial virus infection and novel interventions. Nat Rev Microbiol 21:734–749.

2. Li Y, Wang X, Blau DM, Caballero MT, Feikin DR, Gill CJ, Madhi SA, Omer SB, Simoes EAF, Campbell H, Pariente AB, Bardach D, Bassat Q, Casalegno JS, Chakhunashvili G, Crawford N, Danilenko D, Do LAH, Echavarria M, Gentile A, Gordon A, Heikkinen T, Huang QS, Jullien S, Krishnan A, Lopez EL, Markic J, Mira-Iglesias A, Moore HC, Moyes J, Mwananyanda L, Nokes DJ, Noordeen F, Obodai E, Palani N, Romero C, Salimi V, Satav A, Seo E, Shchomak Z, Singleton R, Stolyarov K, Stoszek SK, von Gottberg A, Wurzel D, Yoshida LM, Yung CF, Zar HJ, Respiratory Virus Global Epidemiology N, Nair H, et al. 2022. Global, regional, and national disease burden estimates of acute lower respiratory infections due to respiratory syncytial virus in children younger than 5 years in 2019: a systematic analysis. Lancet 399:2047–2064.

3. Zhang S, Akmar LZ, Bailey F, Rath BA, Alchikh M, Schweiger B, Lucero MG, Nillos LT, Kyaw MH, Kieffer A, Tong S, Campbell H, Beutels P, Nair H, Investigators R. 2020. Cost of Respiratory Syncytial Virus-Associated Acute Lower Respiratory Infection Management in Young Children at the Regional and Global Level: A Systematic Review and Meta-Analysis. J Infect Dis 222:S680–S687.

4. Pneumonia Etiology Research for Child Health Study G. 2019. Causes of severe pneumonia requiring hospital admission in children without HIV infection from Africa and Asia: the PERCH multi-country case-control study. Lancet doi:10.1016/S0140-6736(19)30721-4.

5. Falsey AR, Hennessey PA, Formica MA, Cox C, Walsh EE. 2005. Respiratory syncytial virus infection in elderly and high-risk adults. N Engl J Med 352:1749–59.

6. Munting A, Manuel O. 2021. Viral infections in lung transplantation. J Thorac Dis 13:6673–6694.

7. Ackerson B, Tseng HF, Sy LS, Solano Z, Slezak J, Luo Y, Fischetti CA, Shinde V. 2019. Severe Morbidity and Mortality Associated With Respiratory Syncytial Virus Versus Influenza Infection in Hospitalized Older Adults. Clin Infect Dis 69:197–203.

8. Korsten K, Adriaenssens N, Coenen S, Butler C, Ravanfar B, Rutter H, Allen J, Falsey A, Pircon JY, Gruselle O, Pavot V, Vernhes C, Balla-Jhagjhoorsingh S, Oner D, Ispas G, Aerssens J, Shinde V, Verheij T, Bont L, Wildenbeest J, investigators R. 2021. Burden of respiratory syncytial virus infection in community-dwelling older adults in Europe (RESCEU): an international prospective cohort study. Eur Respir J 57.

9. Falsey AR, Walsh EE, House S, Vandenijck Y, Ren X, Keim S, Kang D, Peeters P, Witek J, Ispas G. 2021. Risk Factors and Medical Resource Utilization of Respiratory Syncytial Virus, Human Metapneumovirus, and Influenza-Related Hospitalizations in Adults-A Global Study During the 2017-2019 Epidemic Seasons (Hospitalized Acute Respiratory Tract Infection [HARTI] Study). Open Forum Infect Dis 8:ofab491.

10. Heppe-Montero M, Gil-Prieto R, Del Diego Salas J, Hernandez-Barrera V, Gil-de-Miguel A. 2022. Impact of Respiratory Syncytial Virus and Influenza Virus Infection in the Adult Population in Spain between 2012 and 2020. Int J Environ Res Public Health 19.

11. Busack B, Shorr AF. 2022. Going Viral-RSV as the Neglected Adult Respiratory Virus. Pathogens 11.

12. Maggi S, Veronese N, Burgio M, Cammarata G, Ciuppa ME, Ciriminna S, Di Gennaro F, Smith L, Trott M, Dominguez LJ, Giammanco GM, De Grazia S, Costantino C, Vitale F, Barbagallo M. 2022. Rate of Hospitalizations and Mortality of Respiratory Syncytial Virus Infection Compared to Influenza in Older People: A Systematic Review and Meta-Analysis. Vaccines (Basel) 10.

13. Wilson E, Goswami J, Baqui AH, Doreski PA, Perez-Marc G, Zaman K, Monroy J, Duncan CJA, Ujiie M, Ramet M, Perez-Breva L, Falsey AR, Walsh EE, Dhar R, Wilson L, Du J, Ghaswalla P, Kapoor A, Lan L, Mehta S, Mithani R, Panozzo CA, Simorellis AK, Kuter BJ, Schodel F, Huang W, Reuter C, Slobod K, Stoszek SK, Shaw CA, Miller JM, Das R, Chen GL, Conquer RSVSG. 2023. Efficacy and Safety of an mRNA-Based RSV PreF Vaccine in Older Adults. N Engl J Med 389:2233–2244.

14. Mazur NI, Terstappen J, Baral R, Bardaji A, Beutels P, Buchholz UJ, Cohen C, Crowe JE, Jr., Cutland CL, Eckert L, Feikin D, Fitzpatrick T, Fong Y, Graham BS, Heikkinen T, Higgins D, Hirve S, Klugman KP, Kragten-Tabatabaie L, Lemey P, Libster R, Lowensteyn Y, Mejias A, Munoz FM, Munywoki PK, Mwananyanda L, Nair H, Nunes MC, Ramilo O, Richmond P, Ruckwardt TJ, Sande C, Srikantiah P, Thacker N, Waldstein KA, Weinberger D, Wildenbeest J, Wiseman D, Zar HJ, Zambon M, Bont L. 2022. Respiratory syncytial virus prevention within reach: the vaccine and monoclonal antibody landscape. Lancet Infect Dis doi:10.1016/S1473-3099(22)00291-2.

15. Ruckwardt TJ. 2023. The road to approved vaccines for respiratory syncytial virus. NPJ Vaccines 8:138.

16. Afonso CL, Amarasinghe GK, Banyai K, Bao Y, Basler CF, Bavari S, Bejerman N, Blasdell KR, Briand FX, Briese T, Bukreyev A, Calisher CH, Chandran K, Cheng J, Clawson AN, Collins PL, Dietzgen RG, Dolnik O, Domier LL, Durrwald R, Dye JM, Easton AJ, Ebihara H, Farkas SL, Freitas-Astua J, Formenty P, Fouchier RA, Fu Y, Ghedin E, Goodin MM, Hewson R, Horie M, Hyndman TH, Jiang D, Kitajima EW, Kobinger GP, Kondo H, Kurath G, Lamb RA, Lenardon S, Leroy EM, Li CX, Lin XD, Liu L, Longdon B, Marton S, Maisner A, Muhlberger E, Netesov SV, Nowotny N, et al. 2016. Taxonomy of the order Mononegavirales: update 2016. Archives of virology 161:2351–60.

17. Collins PL, Crowe JE. 2007. Respiratory Syncytial Virus and Metapneumovirus. In Fields Virology, pp 1601–1646 Edited by D M Knipe & P M Howley Philadelphia: Lippincott Williams & Wilkins Fifth edition:1601-46.

18. Cao D, Gao Y, Liang B. 2021. Structural Insights into the Respiratory Syncytial Virus RNA Synthesis Complexes. Viruses 13.

19. Garcia J, Garcia-Barreno B, Vivo A, Melero JA. 1993. Cytoplasmic inclusions of respiratory syncytial virus-infected cells: formation of inclusion bodies in transfected cells that coexpress the nucleoprotein, the phosphoprotein, and the 22K protein. Virology 195:243–7.

20. Munday DC, Wu W, Smith N, Fix J, Noton SL, Galloux M, Touzelet O, Armstrong SD, Dawson JM, Aljabr W, Easton AJ, Rameix-Welti MA, de Oliveira AP, Simabuco FM, Ventura AM, Hughes DJ, Barr JN, Fearns R, Digard P, Eleouet JF, Hiscox JA. 2015. Interactome analysis of the human respiratory syncytial virus RNA polymerase complex identifies protein chaperones as important cofactors that promote L-protein stability and RNA synthesis. Journal of virology 89:917–30.

21. Richard CA, Rincheval V, Lassoued S, Fix J, Cardone C, Esneau C, Nekhai S, Galloux M, Rameix-Welti MA, Sizun C, Eleouet JF. 2018. RSV hijacks cellular protein phosphatase 1 to regulate M2-1 phosphorylation and viral transcription. PLoS Pathog 14:e1006920.

22. Wei T, Li D, Marcial D, Khan M, Lin MH, Snape N, Ghildyal R, Harrich D, Spann K. 2014. The eukaryotic elongation factor 1A is critical for genome replication of the paramyxovirus respiratory syncytial virus. PLoS One 9:e114447.

23. Abbas W, Kumar A, Herbein G. 2015. The eEF1A Proteins: At the Crossroads of Oncogenesis, Apoptosis, and Viral Infections. Front Oncol 5:75.

24. Snape N, Li D, Wei T, Jin H, Lor M, Rawle DJ, Spann KM, Harrich D. 2018. The eukaryotic translation elongation factor 1A regulation of actin stress fibers is important for infectious RSV production. Virol J 15:182.

25. Losada A, Munoz-Alonso MJ, Garcia C, Sanchez-Murcia PA, Martinez-Leal JF, Dominguez JM, Lillo MP, Gago F, Galmarini CM. 2016. Translation Elongation Factor eEF1A2 is a Novel Anticancer Target for the Marine Natural Product Plitidepsin. Sci Rep 6:35100.

26. White KM, Rosales R, Yildiz S, Kehrer T, Miorin L, Moreno E, Jangra S, Uccellini MB, Rathnasinghe R, Coughlan L, Martinez-Romero C, Batra J, Rojc A, Bouhaddou M, Fabius JM, Obernier K, Dejosez M, Guillen MJ, Losada A, Aviles P, Schotsaert M, Zwaka T, Vignuzzi M, Shokat KM, Krogan NJ, Garcia-Sastre A. 2021. Plitidepsin has potent preclinical efficacy against SARS-CoV-2 by targeting the host protein eEF1A. Science 371:926–931.

27. Buchholz UJ, Finke S, Conzelmann KK. 1999. Generation of bovine respiratory syncytial virus (BRSV) from cDNA: BRSV NS2 is not essential for virus replication in tissue culture, and the human RSV leader region acts as a functional BRSV genome promoter. J Virol 73:251–9.

28. Rameix-Welti MA, Le Goffic R, Herve PL, Sourimant J, Remot A, Riffault S, Yu Q, Galloux M, Gault E, Eleouet JF. 2014. Visualizing the replication of respiratory syncytial virus in cells and in living mice. Nature communications 5:5104.

29. Bouillier C, Rincheval V, Sitterlin D, Blouquit-Laye S, Desquesnes A, Eleouet JF, Gault E, Rameix-Welti MA. 2019. Generation, Amplification, and Titration of Recombinant Respiratory Syncytial Viruses. J Vis Exp doi:10.3791/59218.

30. Blondot ML, Dubosclard V, Fix J, Lassoued S, Aumont-Nicaise M, Bontems F, Eleouet JF, Sizun C. 2012. Structure and functional analysis of the RNA- and viral phosphoprotein-binding domain of respiratory syncytial virus M2-1 protein. PLoS pathogens 8:e1002734.

31. Sourimant J, Rameix-Welti MA, Gaillard AL, Chevret D, Galloux M, Gault E, Eleouet JF. 2015. Fine mapping and characterization of the L-polymerase-binding domain of the respiratory syncytial virus phosphoprotein. Journal of virology 89:4421–33.

32. Castagne N, Barbier A, Bernard J, Rezaei H, Huet JC, Henry C, Da Costa B, Eleouet JF. 2004. Biochemical characterization of the respiratory syncytial virus P-P and P-N protein complexes and localization of the P protein oligomerization domain. J Gen Virol 85:1643–53.

33. Rolfs Z, Frey BL, Shi X, Kawai Y, Smith LM, Welham NV. 2021. An atlas of protein turnover rates in mouse tissues. Nat Commun 12:6778.

34. Gilman MSA, Liu C, Fung A, Behera I, Jordan P, Rigaux P, Ysebaert N, Tcherniuk S, Sourimant J, Eleouet JF, Sutto-Ortiz P, Decroly E, Roymans D, Jin Z, McLellan JS. 2019. Structure of the Respiratory Syncytial Virus Polymerase Complex. Cell doi:10.1016/j.cell.2019.08.014.

35. Sutto-Ortiz P, Tcherniuk S, Ysebaert N, Abeywickrema P, Noel M, Decombe A, Debart F, Vasseur JJ, Canard B, Roymans D, Rigaux P, Eleouet JF, Decroly E. 2021. The methyltransferase domain of the Respiratory Syncytial Virus L protein catalyzes cap N7 and 2’-O-methylation. PLoS Pathog 17:e1009562.

36. Yu X, Abeywickrema P, Bonneux B, Behera I, Anson B, Jacoby E, Fung A, Adhikary S, Bhaumik A, Carbajo RJ, De Bruyn S, Miller R, Patrick A, Pham Q, Piassek M, Verheyen N, Shareef A, Sutto-Ortiz P, Ysebaert N, Van Vlijmen H, Jonckers THM, Herschke F, McLellan JS, Decroly E, Fearns R, Grosse S, Roymans D, Sharma S, Rigaux P, Jin Z. 2023. Structural and mechanistic insights into the inhibition of respiratory syncytial virus polymerase by a non-nucleoside inhibitor. Commun Biol 6:1074.

37. Morey TM, Esmaeili MA, Duennwald ML, Rylett RJ. 2021. SPAAC Pulse-Chase: A Novel Click Chemistry-Based Method to Determine the Half-Life of Cellular Proteins. Front Cell Dev Biol 9:722560.

38. Carelli JD, Sethofer SG, Smith GA, Miller HR, Simard JL, Merrick WC, Jain RK, Ross NT, Taunton J. 2015. Ternatin and improved synthetic variants kill cancer cells by targeting the elongation factor-1A ternary complex. Elife 4.

39. Oltion K, Carelli JD, Yang T, See SK, Wang HY, Kampmann M, Taunton J. 2023. An E3 ligase network engages GCN1 to promote the degradation of translation factors on stalled ribosomes. Cell 186:346–362 e17.

40. Sasikumar AN, Perez WB, Kinzy TG. 2012. The many roles of the eukaryotic elongation factor 1 complex. Wiley Interdiscip Rev RNA 3:543–55.

41. Li D, Wei T, Abbott CM, Harrich D. 2013. The unexpected roles of eukaryotic translation elongation factors in RNA virus replication and pathogenesis. Microbiol Mol Biol Rev 77:253–66.

42. Mateyak MK, Kinzy TG. 2010. eEF1A: thinking outside the ribosome. J Biol Chem 285:21209–13.

43. Xu B, Liu L, Song G. 2021. Functions and Regulation of Translation Elongation Factors. Front Mol Biosci 8:816398.

44. Negrutskii BS, Shalak VF, Novosylna OV, Porubleva LV, Lozhko DM, El’skaya AV. 2023. The eEF1 family of mammalian translation elongation factors. BBA Adv 3:100067.

45. Lee S, Francoeur AM, Liu S, Wang E. 1992. Tissue-specific expression in mammalian brain, heart, and muscle of S1, a member of the elongation factor-1 alpha gene family. J Biol Chem 267:24064–8.

46. Das T, Mathur M, Gupta AK, Janssen GM, Banerjee AK. 1998. RNA polymerase of vesicular stomatitis virus specifically associates with translation elongation factor-1 alphabetagamma for its activity. Proc Natl Acad Sci U S A 95:1449–54.

47. Qanungo KR, Shaji D, Mathur M, Banerjee AK. 2004. Two RNA polymerase complexes from vesicular stomatitis virus-infected cells that carry out transcription and replication of genome RNA. Proc Natl Acad Sci U S A 101:5952–7.

48. Jimenez PC, Wilke DV, Branco PC, Bauermeister A, Rezende-Teixeira P, Gaudencio SP, Costa-Lotufo LV. 2020. Enriching cancer pharmacology with drugs of marine origin. Br J Pharmacol 177:3–27.

49. Varona JF, Landete P, Lopez-Martin JA, Estrada V, Paredes R, Guisado-Vasco P, Fernandez de Orueta L, Torralba M, Fortun J, Vates R, Barberan J, Clotet B, Ancochea J, Carnevali D, Cabello N, Porras L, Gijon P, Monereo A, Abad D, Zuniga S, Sola I, Rodon J, Vergara-Alert J, Izquierdo-Useros N, Fudio S, Pontes MJ, de Rivas B, Giron de Velasco P, Nieto A, Gomez J, Aviles P, Lubomirov R, Belgrano A, Sopesen B, White KM, Rosales R, Yildiz S, Reuschl AK, Thorne LG, Jolly C, Towers GJ, Zuliani-Alvarez L, Bouhaddou M, Obernier K, McGovern BL, Rodriguez ML, Enjuanes L, Fernandez-Sousa JM, Krogan NJ, Jimeno JM, et al. 2022. Preclinical and randomized phase I studies of plitidepsin in adults hospitalized with COVID-19. Life Sci Alliance 5.

50. Aguareles J, Fernandez PV, Carralon-Gonzalez MM, Izquierdo CF, Marti-Ballesteros EM, Fernandez VP, Sotres-Fernandez G, Garcia-Delangue T, LaPetra RGV, Sanchez-Manzano MD, Gutierrez C, Garcia-Coca M, Carnevali-Ruiz D, Barrena-Puertas R, Luque-Pinilla JM, Lloris R, Luepke-Estefan XE, Lopez-Martin JA, Jimeno JM, Guisado-Vasco P. 2023. Outcomes and clinical characteristics of the compassionate use of plitidepsin for immunocompromised adult patients with COVID-19. Int J Infect Dis 135:12–17.

51. Varona JF, Landete P, Paredes R, Vates R, Torralba M, Guisado-Vasco P, Porras L, Munoz P, Gijon P, Ancochea J, Saiz E, Meira F, Jimeno JM, Lopez-Martin JA, Estrada V. 2023. Plitidepsin in adult patients with COVID-19 requiring hospital admission: A long-term follow-up analysis. Front Cell Infect Microbiol 13:1097809.

52. Landete P, Caliman-Sturdza OA, Lopez-Martin JA, Preotescu L, Luca MC, Kotanidou A, Villares P, Iglesias SP, Guisado-Vasco P, Saiz-Lou EM, Del Carmen Farinas-Alvarez M, de Lucas EM, Perez-Alba E, Cisneros JM, Estrada V, Hidalgo-Tenorio C, Poulakou G, Torralba M, Fortun J, Garcia-Ocana P, Lemaignen A, Marcos-Martin M, Molina M, Paredes R, Perez-Rodriguez MT, Raev D, Ryan P, Meira F, Gomez J, Torres N, Lopez-Mendoza D, Jimeno J, Varona JF. 2024. A Phase III Randomized Controlled Trial of Plitidepsin, a Marine-Derived Compound, in Hospitalized Adults With Moderate COVID-19. Clin Infect Dis 79:910–919.

53. Alonso-Alvarez S, Pardal E, Sanchez-Nieto D, Navarro M, Caballero MD, Mateos MV, Martin A. 2017. Plitidepsin: design, development, and potential place in therapy. Drug Des Devel Ther 11:253–264.

54. Zhang H, Cai J, Yu S, Sun B, Zhang W. 2023. Anticancer Small-Molecule Agents Targeting Eukaryotic Elongation Factor 1A: State of the Art. Int J Mol Sci 24.

55. Papapanou M, Papoutsi E, Giannakas T, Katsaounou P. 2021. Plitidepsin: Mechanisms and Clinical Profile of a Promising Antiviral Agent against COVID-19. J Pers Med 11.

56. Garcia-Fernandez LF, Losada A, Alcaide V, Alvarez AM, Cuadrado A, Gonzalez L, Nakayama K, Nakayama KI, Fernandez-Sousa JM, Munoz A, Sanchez-Puelles JM. 2002. Aplidin induces the mitochondrial apoptotic pathway via oxidative stress-mediated JNK and p38 activation and protein kinase C delta. Oncogene 21:7533–44.

57. Cuadrado A, Garcia-Fernandez LF, Gonzalez L, Suarez Y, Losada A, Alcaide V, Martinez T, Fernandez-Sousa JM, Sanchez-Puelles JM, Munoz A. 2003. Aplidin induces apoptosis in human cancer cells via glutathione depletion and sustained activation of the epidermal growth factor receptor, Src, JNK, and p38 MAPK. J Biol Chem 278:241–50.

58. Losada A, Munoz-Alonso MJ, Martinez-Diez M, Gago F, Dominguez JM, Martinez-Leal JF, Galmarini CM. 2018. Binding of eEF1A2 to the RNA-dependent protein kinase PKR modulates its activity and promotes tumour cell survival. Br J Cancer 119:1410–1420.

59. Losada A, Berlanga JJ, Molina-Guijarro JM, Jimenez-Ruiz A, Gago F, Aviles P, de Haro C, Martinez-Leal JF. 2020. Generation of endoplasmic reticulum stress and inhibition of autophagy by plitidepsin induces proteotoxic apoptosis in cancer cells. Biochem Pharmacol 172:113744.

60. Gordiyenko Y, Llacer JL, Ramakrishnan V. 2019. Structural basis for the inhibition of translation through eIF2alpha phosphorylation. Nat Commun 10:2640.

61. Wong JP, Damania B. 2021. SARS-CoV-2 dependence on host pathways. Science 371:884–885.

62. Molina Molina E, Bech-Serra JJ, Franco-Trepat E, Jarne I, Perez-Zsolt D, Badia R, Riveira-Munoz E, Garcia-Vidal E, Revilla L, Franco S, Tarres-Freixas F, Roca N, Ceada G, Kochanowski K, Raich-Regue D, Erkizia I, Boreika R, Bordoy AE, Soler L, Guil S, Carrillo J, Blanco J, Martinez MA, Paredes R, Losada A, Aviles P, Cuevas C, Vergara-Alert J, Segales J, Clotet B, Ballana E, de la Torre C, Izquierdo-Useros N. 2025. Targeting eEF1A reprograms translation and uncovers broad-spectrum antivirals against cap or m(6)A protein synthesis routes. Nat Commun 16:1087.

63. Losada A, Izquierdo-Useros N, Aviles P, Vergara-Alert J, Latino I, Segales J, Gonzalez SF, Cuevas C, Raich-Regue D, Munoz-Alonso MJ, Perez-Zsolt D, Munoz-Basagoiti J, Rodon J, Chang LA, Warang P, Singh G, Brustolin M, Cantero G, Roca N, Perez M, Bustos-Moran E, White K, Schotsaert M, Garcia-Sastre A. 2024. Plitidepsin as an Immunomodulator against Respiratory Viral Infections. J Immunol 212:1307–1318.

